# A High-Throughput Screen of a Library of Therapeutics Identifies Substrates of P-glycoprotein

**DOI:** 10.1101/528992

**Authors:** Tobie D. Lee, Olivia W. Lee, Kyle R. Brimacombe, Lu Chen, Rajarshi Guha, Sabrina Lusvarghi, Bethilehem G. Tebase, Carleen Klumpp-Thomas, Robert W. Robey, Suresh V. Ambudkar, Min Shen, Michael M. Gottesman, Matthew D. Hall

## Abstract

The ATP-binding cassette transporter P-glycoprotein (P-gp) is known to limit brain penetration of many chemotherapy drugs. Although Food and Drug Administration guidelines require that potential interactions of investigational drugs with P-gp be explored, often this information does not enter into the literature. As such, we developed a high-throughput screen (HTS) to identify substrates of P-gp from a series of chemical libraries, testing a total of 10,804 compounds, most of which have known mechanisms of action. We used the CellTiter-Glo viability assay to test library compounds against parental KB-3-1 human cervical adenocarcinoma cells and the colchicine-selected sub-line KB-8-5-11 that over-expresses P-gp. KB-8-5-11 cells were also tested in the presence of a P-gp inhibitor (tariquidar) to assess reversability of transporter-mediated resistance. Of the tested compounds, a total of 90 P-gp substrates were identified including 55 newly identified P-gp substrates. Substrates were confirmed using an orthogonal killing assay against HEK-293 cells transfected with P-gp. We confirmed that AT7159 (cyclin-dependent kinase inhibitor); AT9283, (Janus kinase 2/3 inhibitor); ispinesib (kinesin spindle protein inhibitor); gedatolisib (PKI-587, phosphoinositide 3-kinase/mammalian target of rampamycin inhibitor); GSK-690693 (AKT inhibitor); and KW-2478 (heat shock protein 90 inhibitor) were substrates, and direct ATPase stimulation was assessed. ABCG2 was also found to confer high levels of resistance to AT9283, GSK-690693 and gedatolisib, while ispinesib, AT7519 and KW-2478 were weaker substrates. Combinations of P-gp substrates and inhibitors were assessed to demonstrate on-target synergistic cell killing. This data will be of use in determining understanding how chemotherapeutic agents will cross the blood-brain barrier.

## Introduction

The ATP-binding cassette (ABC) transporters P-glycoprotein (P-gp, encoded by the *MDR1* gene and later renamed *ABCB1*) and ABCG2 (or breast cancer resistance protein, encoded by the *ABCG2* gene) play major roles in limiting the oral bioavailability of compounds and preventing drug ingress at the blood-brain barrier (BBB) by keeping toxins, drugs, and other compounds out of the brain (Gottesman et al., 2016). Soon after its identification as a drug transporter, P-gp was found to be expressed in the small intestine and colon, liver, pancreas, and kidney (Thiebaut et al., 1987), and pharmacokinetic studies in mice deficient for one of the murine homologs of human *ABCB1, Mdr1a* (renamed *Abcb1a*) demonstrated increased bioavailability of orally-administered taxol compared to wild-type mice (Sparreboom et al., 1997). Likewise, ABCG2 was detected in the small intestine and colon (Fetsch et al., 2006; Maliepaard et al., 2001) and the role of ABCG2 in limiting oral uptake of topotecan was confirmed in mice lacking *Abcg2* expression, the murine homolog of *ABCG2* (Basseville et al., 2016; Jonker et al., 2000).

In addition to being highly expressed in the gastrointestinal tract, in the brush border of renal proximal tubule cells and on the apical surface of hepatocytes (Fetsch et al., 2006; Huls et al., 2008; Thiebaut et al., 1987), both P-gp and ABCG2 are expressed at high levels on the apical side of capillary endothelial cells in the brain (Cooray et al., 2002; Cordon-Cardo et al., 1989; Thiebaut et al., 1987; Thiebaut et al., 1989). The protective role of P-gp was demonstrated in 1994 when Schinkel and colleagues found that deletion of *Abcb1a* in mice resulted in acute sensitivity to the acaricide ivermectin due to a 90-fold increase in brain penetration of the drug (Schinkel et al., 1994). Brain penetration of the P-gp substrate drug vinblastine was increased 20-fold in Abcb1a-deficient mice (Schinkel et al., 1994). Subsequent to the discovery of ABCG2, mice deficient in the two murine homologs of human ABCB1 (*Abcb1a/1b*) and *Abcg2* were generated. The murine models highlighted a compensatory and possibly a cooperative role for the two transporters at the BBB limiting the brain penetration of chemotherapeutic agents, in particular kinase inhibitors (Basseville et al., 2016). In a recent example, 24 h after mice were given an oral dose of the BCR-ABL kinase inhibitor ponatinib, mice lacking *Abcg2* expression had a 2.2-fold increase in brain concentration compared to wild-type mice, while mice lacking *Abcb1a/1b* had a 1.9-fold increase and mice lacking *Abcb1a/1b* and *Abcg2* had a 25.5-fold increase (Kort et al., 2017). The mouse studies highlight not only the protective and complementary role of the transporters at the BBB but also their importance in thwarting effective delivery of chemotherapy to the brain (Robey et al., 2018).

Because transporters affect drug efficacy and pharmacokinetics, it is important to know which compounds are substrates. This can affect decisions on how a drug is administered or whether or not the drug might be effective in the treatment of neurological diseases, and against drug-resistant tumor cells. Although the FDA offers guidelines for determining the interaction of investigational drugs with P-gp and ABCG2 (Lee et al., 2017), often these critical data are not published.

We implemented a systematic screen to identify cytotoxic substrates of P-gp. To do so, we developed high-throughput assays to examine differential cell killing between drug-naïve cancer cell lines and drug-selected, P-gp-overexpressing sublines. We screened several libraries of annotated compounds including the NCATS Pharmaceutical Collection (NPC), a comprehensive collection of all clinically approved drugs, along with small molecules with known mechanisms of action – either as probe small molecules or experimental therapeutics designed to modulate a wide range of targets, including many small molecules developed for oncology indications. Mechanistic annotation of these compounds can provide valuable insight into targets or pathways that appear to have been rendered more sensitive to inhibition in the course of multidrug resistance development. Hits from the primary screen were assessed against HEK cells over-expressing P-gp in the absence and presence of the P-gp inhibitor tariquidar. Top substrates were tested in a cell-killing synergy experiment with the inhibitors tariquidar or elacridar, demonstrating the consistent inhibition of P-gp by inhibitors irrespective of substrate.

## Materials and Methods

### Cell Lines

The HeLa derivative cell line, KB-3-1, and its colchicine-selected, P-gp-overexpressing subline, KB-8-5-11 were maintained in DMEM with 10% FCS and Pen/Strep with glutamine at 37°C in 5% CO2. For KB-8-5-11 cells, colchicine was added to the medium at a concentration of 100 ng/mL. HEK-293 cells transfected with empty vector (pcDNA) or vector containing human *ABCB1* (MDR-19) or *ABCG2* (R-5) have been described previously (Robey et al., 2011) and were maintained in EMEM supplemented with 10% FCS, Pen/Strep and glutamine with 2 mg/ml G418 to select for the expression of the transporter. Cultures were confirmed to be free of mycoplasma infection using the MycoAlert Mycoplasma Detection Kit (Lonza, Walkersville, MD). For the screen, assay medium was identical to culture medium except for KB-8-5-11 where colchicine was excluded from the medium.

### Screening Libraries

The libraries that were used for the screen included the Mechanism Interrogation Plate (MIPE) comprised of 1,912 compounds (Mathews Griner et al., 2014), the NCATS pharmaceutical collection (NPC) comprised of 2816 compounds (Huang et al., 2011), the NCATS Pharmacologically Active Chemical Toolbox (NPACT) (Davis et al., 2016) comprised of 5,099 compounds, and a kinase inhibitor library comprised of 977 compounds, for a total of 10,804 compounds.

### High Throughput Screen (HTS)

All cell lines were plated into 1536-well plates at 500 cells/well in 5 μL media. Compounds were then pinned in dose-response using a 1536-head pin tool (Kalypsis, San Diego, CA) and plates were incubated at 37 °C in 5% CO2 for an additional 72 h. CellTiter-Glo reagent (Promega) was dispensed into the wells, incubated for 5 min and luminescence was read on a ViewLux instrument (Perkin-Elmer). Cytotoxic compounds were defined as those that yielded a curve class of −1.1, −1.2, −2.1, −2.2, −2.3, or −2.4, a maximum response of >50% and an AC50 of ≤ 10μM. Cherry-picked hits from screening analysis were tested with both the KB pair of cell lines, and the pcDNA (empty vector control) and MDR-19 (P-gp overexpressing) pair were tested, in the absence and presence of tariquidar.

Confirmatory three-day cytotoxicity assays were also performed by plating pcDNA, MDR-19 or R-5 cells in 96-well, opaque white plates at a density of 5,000 cells/well and allowing them to attach overnight. Compounds were added at increasing concentrations and incubated with the cells for 72 h after which plates were analyzed using CellTiter-Glo according to the manufacturer’s instructions.

### Synergy Experiments

Synergy screens were performed with a subset of P-gp substrates identified by HTS in combination with the P-gp inhibitors tariquidar or elacridar. Plating of compounds in matrix format using acoustic droplet ejection and numerical characterization of synergy, additivity and/or antagonism were conducted as described previously (Martinez et al., 2016; Mathews Griner et al., 2014). Briefly, compounds were plated as a 10 x 10 dose response combination matrix. Concentration ranges were selected from single agent dose response curves generated from the HTS. Compounds were acoustically dispensed (10 nL/well) using an ATS-100 (EDC Biosystems) onto 1,536-well, white, solid-bottom, TC-treated plates. KB-3-1 and KB-8-5-11 cells were subsequently added to the plates (500 cells/well in 5 μl) and incubated for 72 hr at 37 °C with 5% CO2 under 85% humidity. Cell viability was determined by the addition of 2.5 μL of CellTiter-Glo into to each well. After 15 min incubation at RT, each sample’s luminescence intensity was measured using a ViewLux reader. DMSO (20 nL) and bortezomib (20 nL at 2.3 mM) were used as negative and positive controls respectively. All P-gp substrates listed in Table 1 were tested against the P-gp inhibitors tariquidar and elacridar. Combinations were characterized using the Bliss model and summarized using the DBSumNeg metric.

**Table 1.**
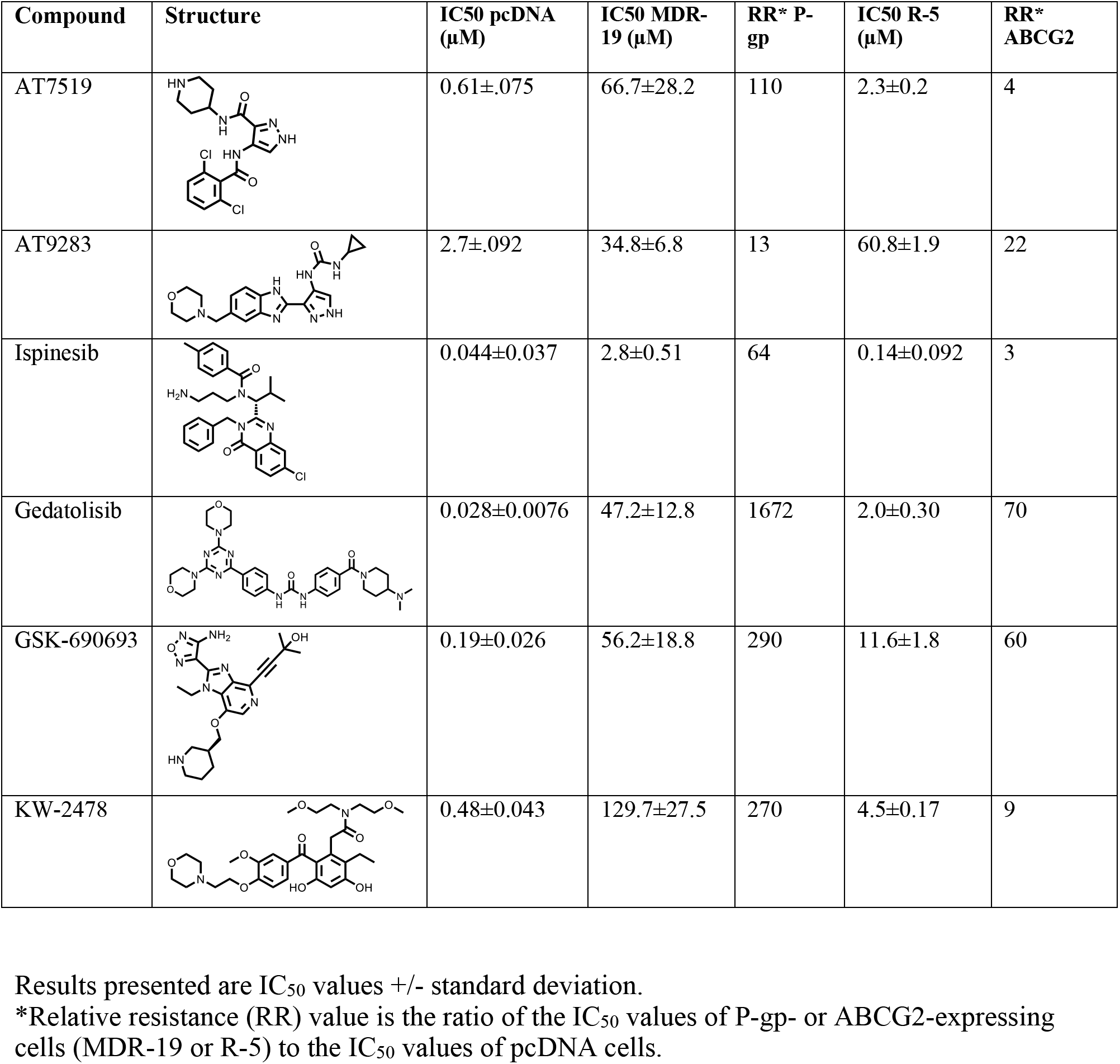
Cross resistance profile for novel P-gp substrates in P-gp- or ABCG2-expressing cells

| Compound | Structure | IC<sub>50</sub> pcDNA (μM) | IC<sub>50</sub> MDR-19 (μM) | RR* P-gp | IC<sub>50</sub> R-5 (μM) | RR* ABCG2 |
| --- | --- | --- | --- | --- | --- | --- |
| AT7519 |  | 0.61±.075 | 66.7±28.2 | 110 | 2.3±0.2 | 4 |
| AT9283 |  | 2.7±.092 | 34.8±6.8 | 13 | 60.8±1.9 | 22 |
| Ispinesib |  | 0.044±0.037 | 2.8±0.51 | 64 | 0.14±0.092 | 3 |
| Gedatolisib |  | 0.028±0.0076 | 47.2±12.8 | 1672 | 2.0±0.30 | 70 |
| GSK-690693 |  | 0.19±0.026 | 56.2±18.8 | 290 | 11.6±1.8 | 60 |
| KW-2478 |  | 0.48±0.043 | 129.7±27.5 | 270 | 4.5±0.17 | 9 |

Results presented are IC<sub>50</sub> values +/- standard deviation.
\*Relative resistance (RR) value is the ratio of the IC<sub>50</sub> values of P-gp- or ABCG2-expressing cells (MDR-19 or R-5) to the IC<sub>50</sub> values of pcDNA cells.

### ATPase Assay

The ATPase assay was performed as described previously (Ambudkar, 1998). Briefly, crude membrane protein (100 μg/ml) was isolated from Hi-Five insect cells expressing human P-gp. The vanadate-sensitive activity was calculated by measuring the end point phosphate release assay in the absence and presence of vanadate. Briefly, solutions containing 4.0 μg of total membrane protein in 100 μL of ATPase assay buffer (50 mM MES-Tris pH 6.8, 50 mM KCl, 5 mM sodium azide, 1 mM EGTA, 1 mM ouabain, 2 mM DTT, 10 mM MgCl2) with 1% DMSO solvent alone (basal activity) or with variable concentrations (0.1, 1 or 10 μM) of the substrates in DMSO were prepared. The tubes were incubated for 3 min at 37°C, after which the reaction was initiated by addition of 5 mM ATP. After 20 min incubation, the reaction was stopped by addition of 2.5% SDS. The amount of inorganic phosphate released was quantified by the colorimetric method, as previously described (Ambudkar, 1998).

### Flow Cytometry

Transport assays were conducted as described previously (Robey et al., 2004). To measure inhibition of P-gp-mediated transport, trypsinized MDR-19 cells were incubated in phenol red-free IMEM supplemented with 10% FCS, Pen/Strep, and glutamine, containing 0.5 μg/mL rhodamine 123 (Sigma-Aldrich, St. Louis MO) in the presence or absence of 25 μM of selected compounds identified by the screen for 30 min at 37°C in 5% CO2. The medium was then removed and replaced with complete medium with or without the compound for an additional 1 h. Valspodar (Apex Biotechnology, Houston, TX) at 3 μg/mL served as a positive control for P-gp inhibition. For ABCG2-mediated transport, R-5 cells were incubated in a similar fashion except 5 μM pheophorbide A (PhA, Frontier Scientific, Logan, UT) was used as the substrate and 10 μM fumitremorgin C (FTC, synthesized by the NIH Chemical Biology Laboratory, Bethesda, MD) served as the positive control for ABCG2 inhibition. Samples were analyzed on a FACSCanto II flow cytometer (BD Biosciences, San Jose, CA) in which rhodamine fluorescence was detected with a 488-nm argon laser and a 530-nm bandpass filter and PhA was detected using a 635-nm red diode laser and a 670-nm filter. At least 10,000 events were collected for each sample.

## Results

### High-throughput Screen to Identify P-gp Substrates

To identify novel substrates of P-gp, we used the CellTiter-Glo luminescent cell viability assay to test library compounds against three cell conditions: (1) the parental KB-3-1 human cervical adenocarcinoma cell line (a HeLa clone); (2) the drug-resistant subline KB-8-5-11 that over-expresses P-glycoprotein, and (3) the KB-8-5-11 cell line in the presence of the P-gp inhibitor tariquidar (which should restore sensitivity to P-gp substrates). Overall, 10,804 compounds were tested from across four annotated small molecule libraries. The NCATS Pharmaceutical Collection (NPC) is a library of compounds approved for use by the Food and Drug Administration and related agencies in foreign countries. The NCATS Pharmacologically Active Chemical Toolbox (NPACT) library contains pre-clinical and probe compounds from across disease areas. The kinase inhibitor library contains almost 1,000 small molecule inhibitors of kinases, with known mechanisms of action. The Mechanism Interrogation PlatE (MIPE) library contains oncology-focused compounds with known mechanisms of action.

The screen was designed based on the fundamental principle of P-gp-mediated drug resistance: substrates which kill or inhibit growth of KB-3-1 were expected to demonstrate reduced efficacy against the KB-8-5-11 cells due to drug efflux by P-gp. P-gp-specific efflux could be demonstrated by sensitization of KB-8-5-11 cells in the presence of the P-gp inhibitor tariquidar. As a cell viability assay was utilized for this screen, a limitation of the screen is that only cytotoxic or cytostatic P-gp substrates could be identified. Of 10,804 compounds tested, 1,362 compounds demonstrated cytotoxicity towards the KB-3-1 cells, of which 90 compounds were identified as putative P-gp substrates (Fig. 1A). In total, 13% of all compounds tested demonstrated cytotoxicity towards the parental KB-3-1 cell line. Among the four libraries, the kinase inhibitor library (30%) and oncology-focused MIPE library (21%) contained the greatest proportion of cytotoxic compounds, followed by the NPACT library (10%) (Fig. 1B). As one might anticipate, the NPC library of therapeutic agents contained the lowest proportion of cytotoxic compounds (5%).

**Fig. 1.**
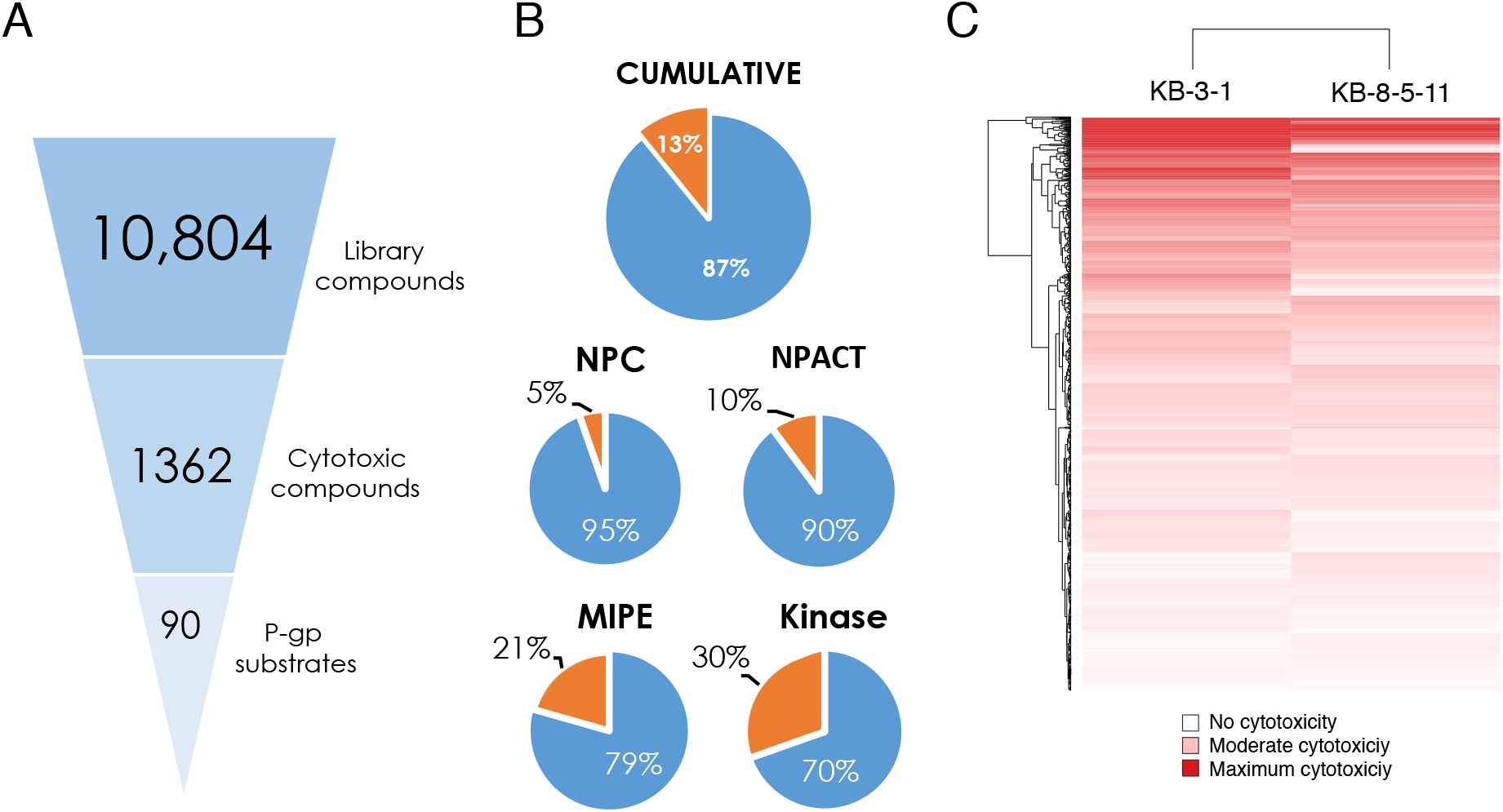
Overview of high-throughput screening data. (A) Hit triage from HTS library screen (10,804 compounds), to compounds that are cytotoxic towards parental KB-3-1 cells (1,362 compounds) to identified P-gp substrates (90 compounds). (B) Summary of percentage of cytotoxic compounds, cumulative and represented in each library screened. (C) Area under the curve (AUC) heatmap of compound activity for screen hits, where red intensity represents magnitude of AUC: strong red is strongest cytotoxicity, and white represents no cytotoxicity.

A comparison of the global response of the two cell lines to the library compounds was undertaken by assessing the area-under-the-curve (AUC) for each compound against each cell line (Fig. 1C). The AUC of the dose-response curve ensures both efficacy (magnitude of cell killing) and potency (concentration that elicits cell killing) are accounted for in the analysis of activity (Fig. 1A). The KB-3-1 cell line was more sensitive than the KB-8-5-11 cell line for a number of compounds, shown in Fig. 1C, where darker red correlates with greater sensitivity to a given compound, while lighter colors correlate with more resistance. This is consistent with the multidrug-resistant nature of the KB-8-5-11 cell line.

To pinpoint substrates identified by HTS, the difference in AUC between KB-3-1 and KB-8-5-11 cells was determined (ΔAUC1, Fig. 2A). As an example, the HTS dose-response curves for vincristine (a known P-gp substrate) are displayed in Fig. 2A for all three conditions screened. As cell killing assays involve loss of signal, data were analyzed from 0% (positive control signal) to −100% (complete cell killing/growth inhibition). The data represented in Fig. 2A is usual for HTS data analysis, but not normal for displaying cell killing data – elsewhere in this manuscript, data are displayed as is traditional for cell killing assays with 100% as control cell viability and 0% as total cell death. The AUC for the sensitive KB-3-1 cell line was determined for each compound (gray shading). The AUC for the resistant KB-8-5-11 cell line was also determined for each compound (red shading). The difference (delta) between the AUC for each cell line was determined (termed ΔAUC1), and the greater the magnitude of ΔAUC1, the stronger the substrate effect of P-gp. This strategy is distinct from the commonly applied method used to discern P-gp substrates by comparing the IC50 values derived from dose-response curves. The difference between KB-8-5-11 cells with and without the P-gp inhibitor tariquidar (1 μM) was also assessed in order to confirm P-gp substrates (termed ΔAUC2). P-gp primary high-throughput screening data for KB-3-1 and KB-8-5-11 cell lines against NPC, NPACT MIPE and Kinase libraries was deposited in PubChem with AIDs 1346986 and 1346987, respectively. Assay data can be accessed via the following links: https://pubchem.ncbi.nlm.nih.gov/assay/assay.cgi?aid=1346986 for the KB-3-1 cell line, and https://pubchem.ncbi.nlm.nih.gov/assay/assay.cgi?aid=1346987 for the KB-8-5-11 cell line.

**Fig. 2.**
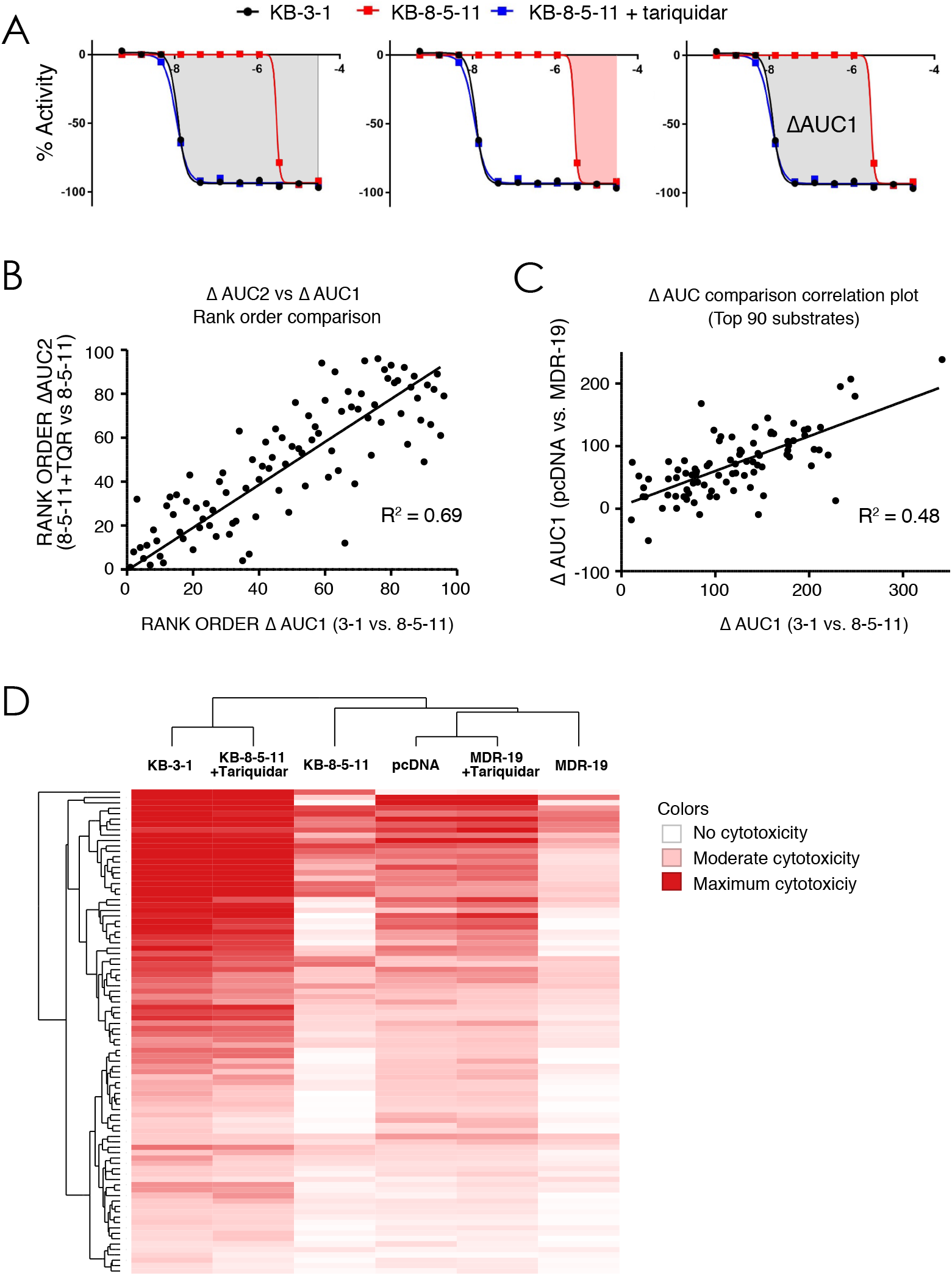
Summary of P-gp substrate analysis. (A) Summary of data analysis. For each compound, data is displayed as ‘loss of signal from baseline’, where −100% represents total cell killing. KB-3-1 cells are sensitive (black), KB-8-5-11 cells are resistant (red) but sensitized when P-gp inhibitor tariquidar is added (blue), top left. To perform HTS data analysis, AUC is determined for parent line (top right, grey) and resistant line (bottom left, pink). The difference in AUC (termed ΔAUC1) is used to identify putative substrates. ΔAUC1 was calculated by difference between resistant cells in the absence and presence of tariquidar. Comparison of the AUC for substrates calculated using ΔAUC1 and ΔAUC2, assessing (B) rank order of substrates (where largest ΔAUC is strongest substrate) and (C) orthogonal data generated testing P-gp substrates against HEK cells. (D) Unsupervised clustering of compound activity against parent and resistant lines (with and without tariquidar), where strong red represents greatest cytoxicity..

Following plating of hits and re-testing for confirmation, a total of 90 P-gp substrates were identified (Supplementary Table 1) based on a ΔAUC1 cut-off value of 50. Comparison of identified substrates with the literature revealed 35 known P-gp substrates were identified in the screen, and 55 new substrates were identified. Comparison of ΔAUC1 and ΔAUC2 for all substrates revealed a strong correlation between parental cells and inhibited KB-8-5-11 cells in which P-gp was inhibited with tariquidar (Fig. 2B). To confirm that the putative P-gp substrates were not due to cell line-specific alterations, an orthogonal assay testing all hits against HEK 293 human embryonic kidney cells stably transfected with either plasmid control (pcDNA) or a plasmid expressing *ABCB1* (MDR-19), was conducted and and ΔAUC1 was calculated. A correlation was demonstrated between the KB-3-1/KB-8-5-11 ΔAUC1 and the pcDNA/MDR-19 ΔAUC1 (Fig. 2C). Unsupervised clustering of global cell response to the 90 substrates identified for the three KB and three HEK conditions (parent, resistant, resistant with tariquidar) demonstrated a consistent pattern, with the KB-8-5-11 and MDR-19 cell lines less sensitive to compounds compared to their parental partners, and the parent lines clustering with the resistant cell lines in the presence of tariquidar (Fig. 2D).

### Assessment of P-gp Substrates

To confirm that the HTS assay and analysis strategy identifies substrates, we assessed the cell-killing activity of three known substrates included in the library (target in brackets): paclitaxel [tubulin] (Greenberger et al., 1988), vincristine [tubulin] (Horton et al., 1987), and mithramycin [RNA synthesis] (Biedler and Riehm, 1970) (Fig. 3A-C, respectively). Examples of the dose-response curves for three newly identified substrates are PKI-402 [PI3K/mTOR](Dehnhardt et al., 2010), CB300919 [NAMPT](Bavetsias et al., 2002), and PHA-793887 [CDK1/2/4](Brasca et al., 2010) (Fig. 3D-F, respectively). In each case, a strong difference in cell killing between parent and resistant cells was found.

**Fig. 3.**
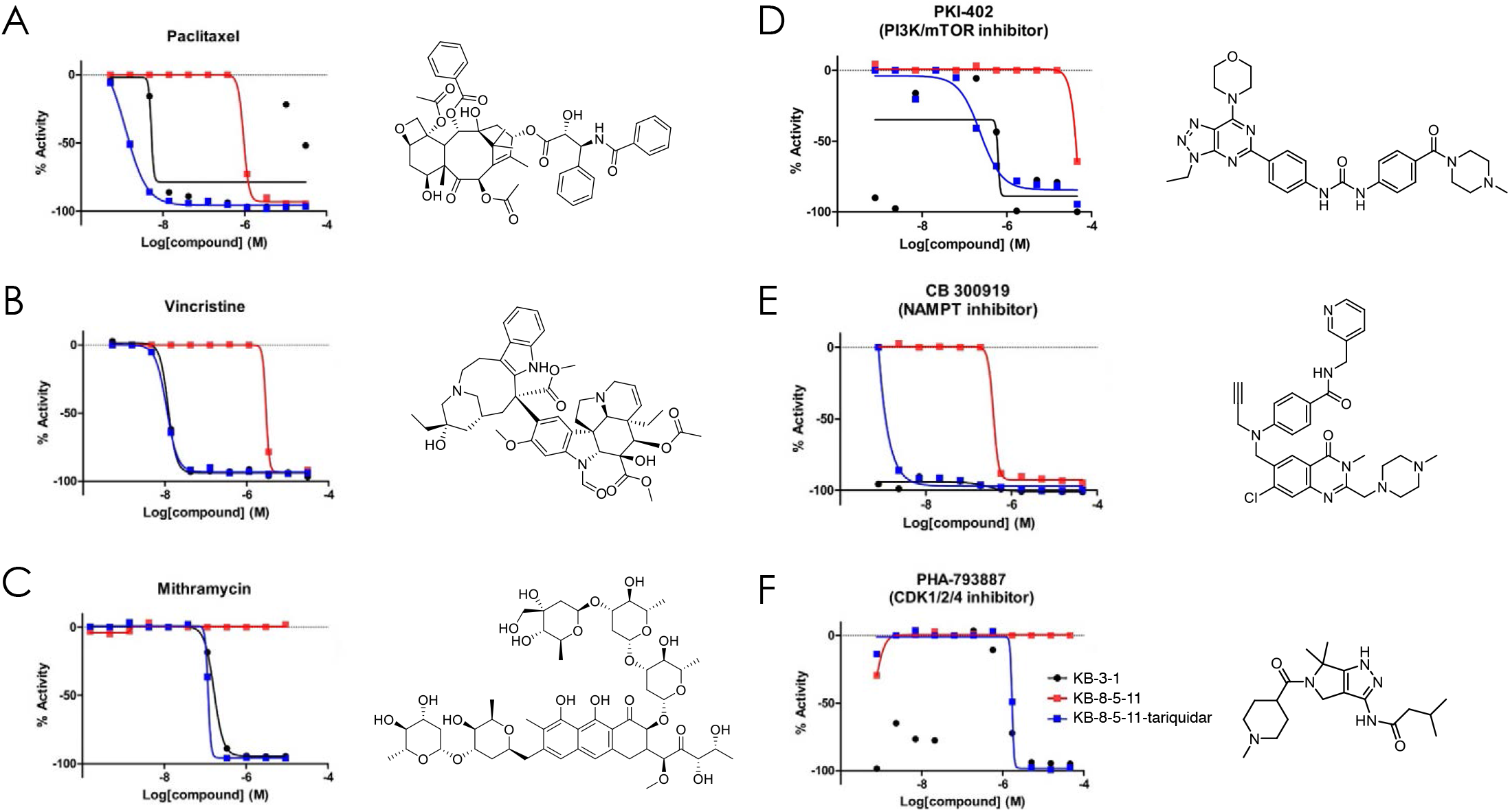
Representative examples of known (A-C) and previously unidentified (D-F) substrates of P-gp.

To confirm that the newly identified substrates were indeed P-gp substrates, we performed confirmatory cytotoxicity assays in the laboratory using pcDNA and MDR-19 cells. We selected 6 commercially available, newly-identified P-gp substrates AT7159, a cyclin-dependent kinase (CDK) inhibitor; AT9283, a JAK2/3 inhibitor; ispinesib, a kinesin spindle protein inhibitor; gedatolisib (PKI-587), a PI3K/mTOR inhibitor; GSK-690693, an AKT inhibitor; and KW-2478, an HSP90 inhibitor. In addition, we examined the ability of ABCG2 to confer resistance to the compounds using ABCG2-overexpressing R-5 cells. As shown in Table 1, all of the compounds were confirmed to be P-gp substrates, with P-gp expression conferring less resistance to AT9283 (13-fold), but conferring very high levels of resistance to gedatolisib (1671-fold). All of the compounds were also found to be ABCG2 substrates to varying degrees. In the case of AT9283, GSK-690693 and gedatolisib, ABCG2 conferred relatively high levels of resistance, while ispinesib, AT7519 and KW-2478 were preferentially transported by P-gp.

### Effect of Some Newly-Identified Substrates on ATPase Activity of P-gp

The ability of substrates to stimulate the ATPase activity of P-gp was assessed. Some kinase inhibitors that are P-gp substrates have been shown to significantly stimulate the ATPase activity of P-gp, while others do not (Hegedus et al., 2009). We omitted AT9283 from this assay, as P-gp conferred the lowest levels of resistance to this compound. The vanadate-sensitive ATPase activity of P-gp in the presence of 0.1, 1 or 10 μM of each compound compared with the basal activity (activity in the presence of 1% DMSO). As observed in Fig. 4A-E, we found that ispinesib stimulated ATPase activity in a concentration-dependent manner. Ispinesib stimulated the ATPase activity by >2.6 fold at concentrations greater than 1 μM (1.6 fold at 0.1 μM). KW-2478 was also capable of stimulating the ATPase activity, but only at higher concentrations (10 μM) and only up to 1.4-fold. The rest of the compounds did not affect the ATPase activity in the range of concentrations tested. Of note, P-gp conferred the highest levels of resistance to gedatolisib in our studies but gedatolisib did not cause an increase in ATPase activity.

**Fig. 4.**
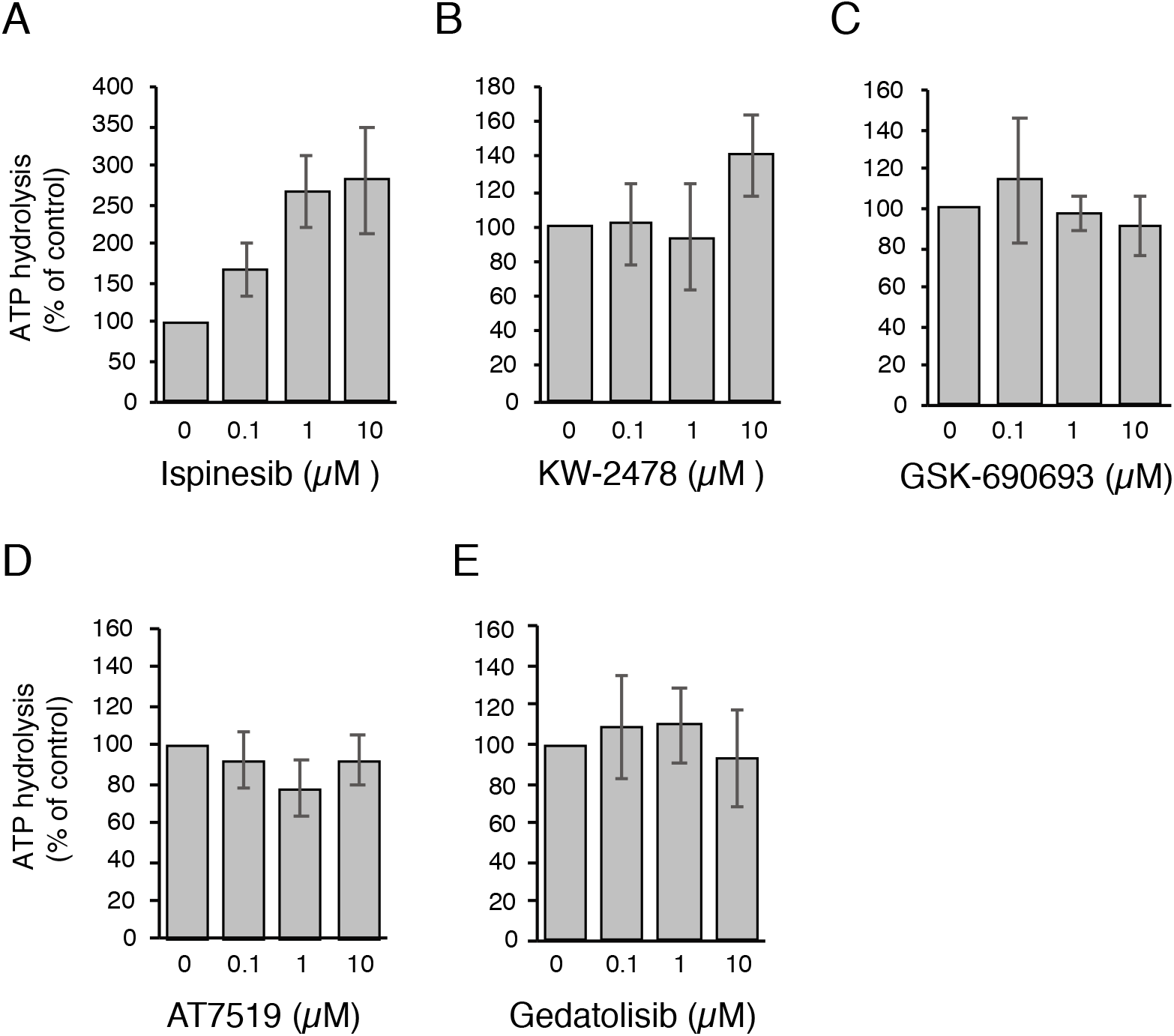
Effects of novel P-gp substrates on ATPase activity. The vanadate-sensitive activity of Pgp was determined as outlined in Materials and Methods. Basal P-gp ATPase activity was compared to activity in the presence of 0.1, 1 or 10 μM of the substrates (A) ispinesib, (B) KW-2478, (C) GSK-690693, (D) AT7519 or (E) gedatolisib. Graphs depict average values from three independent experiments (error bars +/− SD).

### P-gp Inhibitors Synergize With Substrates

P-gp substrates were identified during the HTS in part by co-treating P-gp-expressing cells with the inhibitor tariquidar at a concentration (1 μM) shown to fully inhibit P-gp, to demonstrate P-gp-specific resistance. To explore the nature of inhibitor-mediated sensitization of substrates, we assessed 10 x 10 combinations of tariquidar or elacridar with 17 P-gp substrates identified from the screen (Table 2). Viability was again measured using CellTiter-Glo, and the Bliss independence model was used to characterize the presence or absence of synergy for each combination, where negative ΔBliss represents synergy, and positive ΔBliss represents antagonism.. We hypothesized that inhibitors should antagonize P-gp transport of substrates in P-gp-expressing cells, and that this approach could be used to compare the efficacy of P-gp inhibitors in combination with a range of substrates. This approach has not been previously adopted for studying ABC transporter inhibitors.

**Table 2.**
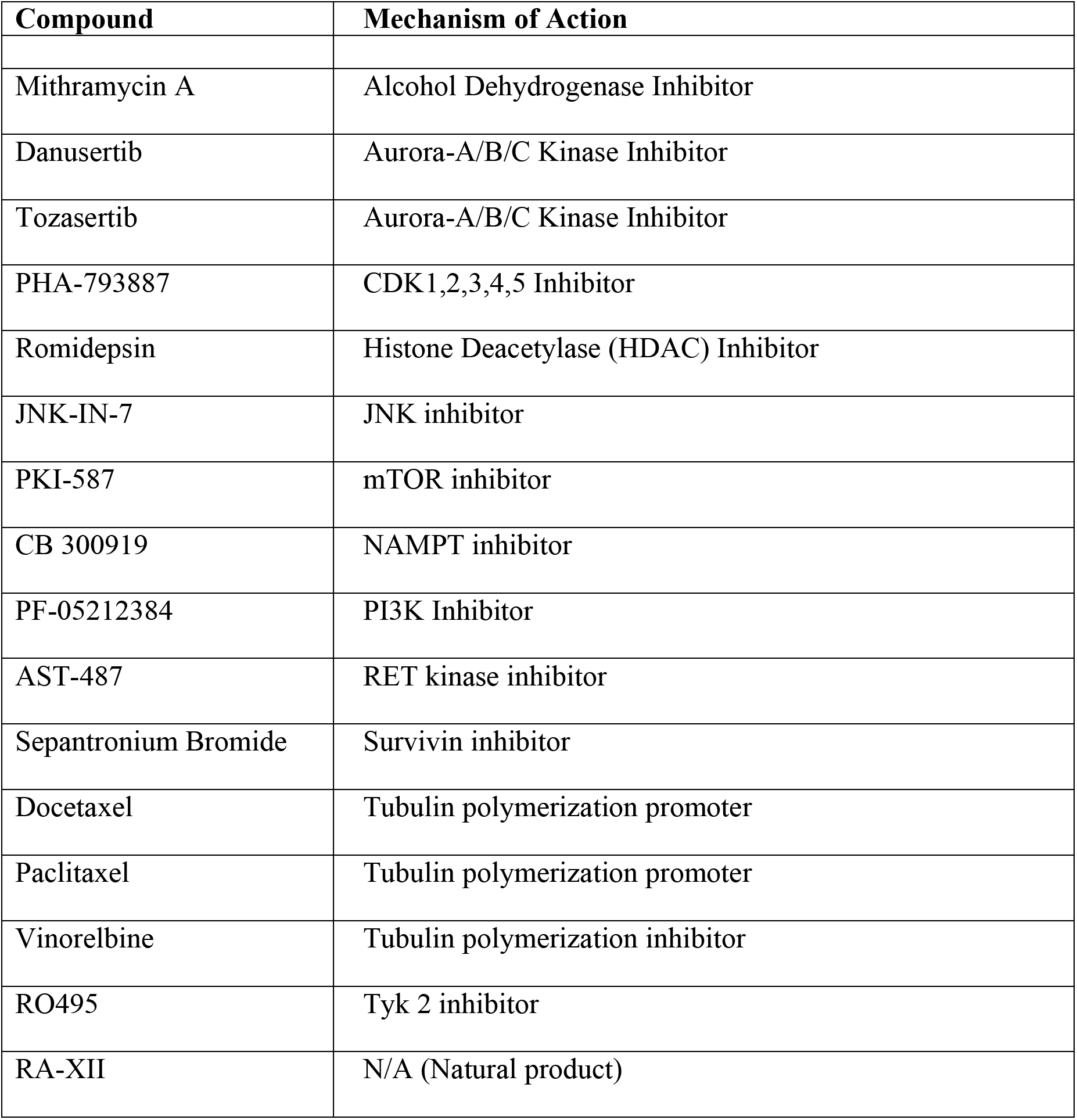
P-gp substrates tested in combination with elacridar and tariquidar

The effect of both tariquidar and elacridar across all substrates was uniform. Paclitaxel is shown as an example. Parental KB-3-1 cells were sensitive to paclitaxel (black = 100% viable, red = 0% viable), and this sensitivity was unaffected by addition of elacridar up to a concentration of 20 μM (Fig. 5a). This is exemplified by the absence of any strong synergy (red) or antagonism (blue) (Fig. 5a), and in paclitaxel dose-response curves extracted from the 10×10 block (Fig. 5b). This relationship was observed for both inhibitors in combination with every Pgp substrate tested against parental cells. Elacridar and tariquidar alone had no effect on cell viability. The 10 x 10 blocks for parental KB-3-1 cell are available at https://tripod.nih.gov/matrix-client/rest/matrix/blocks/8067/table.

**Fig. 5.**
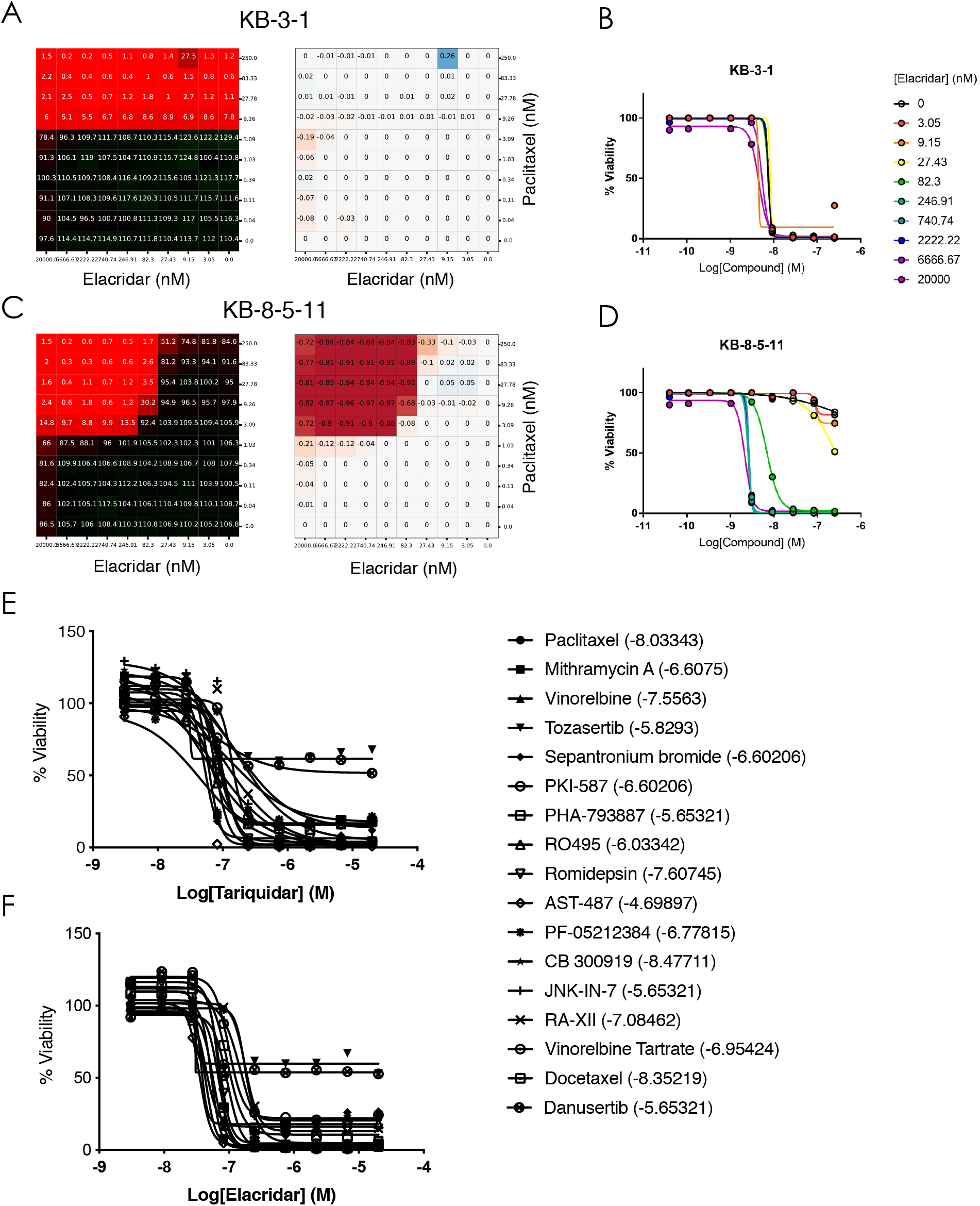
Substrate synergy with P-gp inhibitors. Sample combination of elacridar and paclitaxel tested in 10×10 dose-response matrices with KB-3-1 (A) and KB-8-3-11 (B). Left, percent response of cell viability where red = cell death. Middle, magenta = synergy. Rightdose-response curves extracted from synergy blocks for paclitaxel with increasing concentration of elacridar. Dose-response curves for (E) tariquidar and (F) elacridar for all substrates tested (listed in Table 2).

In contrast, the P-gp substrates demonstrated maximal synergy in combination with elacridar and tariquidar in P-gp-expressing KB-8-5-11 cells. The combination of paclitaxel and elacridar is shown as an example (Fig. 5c, 5d). In the absence of elacridar, paclitaxel demonstrated minimal cytotoxicity towards KB-8-5-11 cells, but addition of elacridar sensitized the cells to paclitaxel, with near-maximal effects at 82 nM and higher. Dose-response curves from the 10 x 10 block demonstrate the sensitization of KB-8-5-11 cells to paclitaxel with increasing elacridar concentration (Fig. 5c), and this sensitization was accompanied by maximal synergy. Synergy was observed for both inhibitors in combination with every P-gp substrate tested against P-gp-expressing KB-8-5-11 cells. The 10 x 10 blocks for P-gp-expressing KB-8-5-11 cell are available at https://tripod.nih.gov/matrix-client/rest/matrix/blocks/8069/table.

To ascertain the inhibitory potency of tariquidar and elacridar, dose-response curves were extracted from the 10 x 10 block of each inhibitor against each compound. 15 of 17 compounds elicited near-complete cell killing, the exceptions being the Aurora kinase inhibitors tozasertib and danusertib, which had a maximal efficacy of approximately 50% (Fig. 5e, 5f). Tariquidar (Fig. 5e) and elacridar (Fig. 5f) both achieved near-complete inhibition of all compounds (maximal cell killing) at 247 nM, and the average IC50 against fifteen diverse substrates were 100 ± 49 nM (cotreated with tariquidar) and 84 ± 48 nM (cotreated with elacridar), suggesting inhibition of P-gp irrespective of the substrate. IC50s for inhibition of compounds were consistent with values in the literature, though there are no studies comparing inhibitors and testing them against a large number of substrates.

### Newly-Identified P-gp Substrates Inhibit P-gp- and ABCG2-Mediated Transport

Many P-gp substrates have also been found to inhibit P-gp-mediated transport at relatively high concentrations and this is particularly true for kinase inhibitors (Durmus et al., 2015). We next assessed the ability of the six compounds identified earlier to inhibit P-gp-mediated rhodamine 123 transport or ABCG2-mediated pheophorbide A transport from MDR-19 or R5 cells, respectively, as shown in Fig. 6. At a concentration of 25 μM, only ispinesib was found to appreciably inhibit P-gp-mediated rhodamine transport and was nearly as effective as 3 μM valspodar. Ispinesib was also the only compound found to appreciably inhibit pheophorbide A transport, although not as well as fumitremorgin C, which served as a positive control for ABCG2 inhibition. In agreement with previous reports, we find that some kinase inhibitors are substrates of transporters at low concentrations, but then act as inhibitors at higher concentrations.

**Fig. 6.**
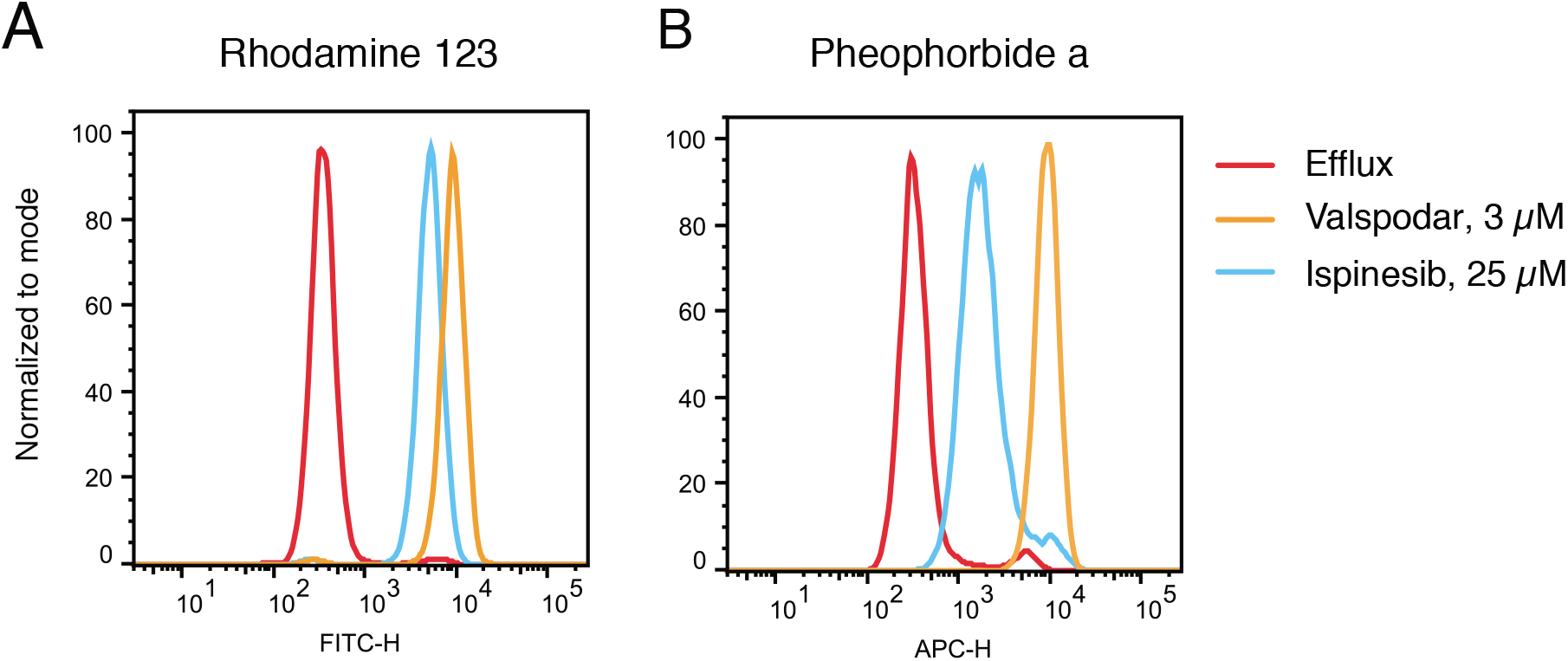
Ispinesib inhibits P-gp- and ABCG2-mediated transport. P-gp overexpressing MDR-19 cells or ABCG2 overexpressing R-5 cells were incubated with 0.5 μg/ml rhodamine 123 or 5 μM pheophorbide A, respectively, in the absence or presence a specific inhibitor (3 μM valspodar for P-gp and 10 μM FTC for ABCG2) or 25 μM ispinesib for 30 min after which the medium was remove and replaced with substate-free medium continuing without or with the inhibitor. Cells incubated with substrates alone are shown by the red histogram, cells incubated with the substrate and specific inhibitor are shown by the orange histogram and cells incubated with the substrate and ispinesib are shown by the blue histogram. Results from one of two experiments are shown.

## Discussion

P-gp and ABCG2 are known to play a role in the disposition of many toxins by limiting oral bioavailability, increasing excretion, and limiting brain penetration (Robey et al., 2018). While many targeted therapies have been developed, often it is unclear what role transporters might play in their disposition and how they might affect therapy. We therefore developed a high-throughput screening assay to identify novel substrates of P-gp. Using KB-3-1 cells that do not express P-gp and their P-gp-overexpressing KB-8-5-11 counterpart, we identified 55 novel substrates of P-gp that were confirmed in a second pair of cell lines both in the primary screen and secondary assays. These data will be valuable to researchers who are seeking novel treatments for brain cancers or metastases, as compounds that are transported by P-gp will most likely not cross the blood-brain barrier, as shown by mouse knockout models (Robey et al., 2018).

Previous screens have used alternative methods to identify novel P-gp substrates. The NCI-60 drug screen cell line set was previously used to identify P-gp substrates based on measuring rhodamine 123 transport (Lee et al., 1994) or *ABCB1* gene expression data (Alvarez et al., 1995; Szakacs et al., 2004) in the 60 cell lines of the screen and comparing that to drug sensitivity profiles. Cell lines with higher levels of P-gp expression were found to correlate with decreased sensitivity to substrate drugs. This method was successful due to the relatively high level of variation of P-gp expression in the lines of the screen. In the case of ABCC1 or ABCG2, with cells expressing much lower levels of the transporters, the screen was less successful and often did not identify known substrates (Alvarez et al., 1998; Deeken et al., 2009). More recently our group developed an assay based on a dual-fluorescent system, in which sensitive cells (OVCAR8) were transduced to express DsRed red fluorescent protein and P-gp overexpressing cells (NCI/ADR-RES) expressed enhanced green fluorescent protein (Brimacombe et al., 2009).

The present study, and those referenced above, rely on P-gp to protect cells from cell death. While appropriate for a study such as this examining cancer drug resistance, a limitation of this approach is that non-toxic compounds cannot be studied. Strategies for studying non-toxic substrates have been explored, and rely on either direct monitoring of the test compounds (for example, radioactivity or analytical detection), or interference with the efflux of a fluorescent substrate. The primary example of the latter comes from Sklar and co-workers, who have reported a number of screens using flow cytometry to identify inhibitors of ABC transporters (Ivnitski-Steele et al., 2008; Strouse et al., 2013a; Strouse et al., 2013b; Winter et al., 2008). A Pfizer study reported a correlation between MDCK cells transfected with mouse (Mdr1a) and human (MDR1) P-gp for 3300 compounds, using LC-MS for analytical quantitation of each compound, although the compounds themselves were not disclosed (Feng et al., 2008). Further work is needed to tabulate all pharmacologically active drugs that are P-gp substrates.

Of course, high-throughput screening methods do have limitations. First, the compounds must be toxic and/or cause cell cycle arrest so that differences between treated and untreated cells can be detected by the CellTiter Glo assay. Thus, the assay will not detect all substrates. Additionally, the ability of a drug to inhibit growth or elicit toxicity often depends on the choice of cell line model. For example, while mutant BRAF inhibitors were among the compounds tested, none emerged as potential substrates despite the fact that some of them such as vemurafenib and dabrafenib were reported to be substrates of P-gp (Mittapalli et al., 2013; Mittapalli et al., 2012). This is not unexpected, as the cell line model systems we used did not harbor a mutant BRAF gene. Among the substrates identified, there were no MEK inhibitors, although trametinib and cobimetinib are both known to be P-gp substrates (Choo et al., 2014; de Gooijer et al., 2017). This is most likely due to the fact that the cell line models used did not harbor mutations in BRAF or Ras. Therefore, while this assay did identify several new substrates of P-gp, it does not represent a definitive way to determine if a compound is a P-gp substrate.

Synergy is an important concept in combination chemotherapy, but it is not often discussed in the context of transport inhibitors. Utilizing the Bliss calculation, we demonstrated the profound synergy that inhibition of P-gp can produce in combination with avid P-gp substrates. An advantage of the 10 x 10 combination grid is the ability to readily examine the effects of inhibitors on P-gp substrates. After the initial screen, the ability of the inhibitors tariquidar and elacridar were tested in increasing concentrations with the 17 substrates identified by the screen. Both inhibitors were found to synergize with the substrates, and to act at a consistent concentration in combination with all compounds tested. The sensitivity of the synergy screening approach suggests that it may be an alternative strategy to identify P-gp substrates, and could potentially be used to detect non-cytotoxic substrates that are also competitive inhibitors.

In conclusion, we have developed a high-throughput screen to identify substrates of P-gp. We identified 55 novel substrates, among them targeted therapies that have yet to be developed clinically. We also have demonstrated that our method can be easily used to confirm the action of proposed P-gp inhibitors by sensitizing P-gp-overexpressing cells to numerous substrates. Future studies will focus on translating these techniques to the identification of substrates of other ABC transporters.

## Acknowledgements

We appreciate the technical assistance of George Leiman. The content of this publication does not necessarily reflect the views or policies of the Department of Health and Human Services, nor does mention of trade names, commercial products, or organizations imply endorsement by the U.S. Government.

## Authorship Contributions

Participated in research design: TDL, OWL, KRB, RWR, SVA, MMG, MDH Conducted experiments: TDL, OWL, LC, RG, SL, BGT, RWR Performed data analysis: TDL, OWL, LC, RG, SL, RWR, SVA, MS Wrote or contributed to the writing of the manuscript: TDL, SL, RWR, MDH

## Non-standard abbreviations

P-gp: P-glycoprotein
BBB: blood-brain barrier
HTS: high-throughput screen
ABCB1: ATP-binding cassette family member B1
ABCG2: ATP-binding cassette family member G2
NCATS: National Center for Advancing Translational Science
NPC: NCATS Pharmaceutical Collection
MIPE: Mechanism Interrogation Plate
NPACT: NCATS Pharmacologically Active Chemical Toolbox
PhA: pheophorbide A
FTC: Fumitremorgin C
AUC: area under the curve
NAMPT: nicotinamide phosphoribosyltransferase
CDK: cyclin-dependent kinase
PI3K: phosphoinositide-3 kinase
mTOR: mammalian target of rapamycin
JAK: Janus kinase

